# Optimizing Trait Predictability in Hybrid Rice Using Superior Prediction Models and Selective Omic Datasets

**DOI:** 10.1101/261263

**Authors:** Shibo Wang, Julong Wei, Ruidong Li, Han Qu, Weibo Xie, Zhenyu Jia

**Author notes:** These authors contributed equally (Shibo Wang, Julong Wei).

## Abstract

Hybrid breeding has dramatically boosted yield and its stability in rice. Genomic prediction further benefits rice breeding by increasing selection intensity and accelerating breeding cycles. With the rapid advancement of technology, other omic data, such as metabolomic data and transcriptomic data, are readily available for predicting genetic values (or breeding values) for agronomically important traits. In the current study, we searched for the best prediction strategy for four traits (yield, 1000 grain weight, number of grains per panicle and number of tillers per plant) of hybrid rice by evaluating all possible combinations of omic datasets with different prediction methods. We conclude that, in rice, the predictions using the combination of genomic and metabolomic data generally produce better results than single-omics predictions or predictions based on other combined omic data. Inclusion of transcriptomic data does not improve predictability possibly because transcriptome does not provide more information for the trait than the sum of genome and metabolome; rather, the computational complexity is substantially increased if transcriptomic data is included in the models. Best linear unbiased prediction (BLUP) appears to be the most efficient prediction method compared to the other commonly used approaches, including LASSO, SSVS, SVM-RBF, SVP-POLY and PLS. Our study has provided a guideline for selection of hybrid rice in terms of which types of omic datasets and which method should be used to achieve higher trait predictability.

## Introduction

Rice, which is enriched with complex carbohydrates, vitamins, minerals, and fiber, is the main staple food for a large segment of the world population. Heterosis, referred to the superior performance of hybrids relative to their parents, has been reported as a major contributor to the increased productivity in rice (Jones, 1926; Virmani et al., 1981). Only a small number of desirable hybrids can be selected through a large number of crosses in a traditional rice breeding program which is labor intensive and time consuming (Collard and Mackill 2008; Spindel et al. 2015). Marker-assisted selection (MAS) has been used to facilitate rice breeding (Chen et al. 2000; Chen et al. 2001; Zhou et al. 2003), leading to genetic improvement and reduced generation time. Quantitative trait loci (QTL) mapping is often used to identify DNA markers for breeding if these markers are in linkage disequilibrium (LD) with the genetic determinant of traits (Asins 2002). Genomic selection (Hayes and Goddard 2001) is a special form of MAS in which all markers on the genome are used for predicting expected breeding values (EBVs) for rice hybrids. A training set is used to build a genomic selection model which can be applied to an independent set for prediction of EBVs if this set share similar genetic architecture with the training set. Genomic selection models are often evaluated by trait predictability, a measurement of prediction accuracy that is calculated through cross validation (Riedelsheimer et al. 2012). A primary goal of genomic selection modelling is to optimize the trait predictability, which is defined as the squared correlation between the observed and the predicted phenotypic values.

In addition to genomic data, the rapid advancement of technology generates other types of omic datasets, such as transcriptomic data, proteomic data, and metabolomic data. An integrated analysis of these omic datasets may advance our knowledge of the underlying genetic and biochemical basis for agronomic traits. For example, the joint analysis of transcriptomic data and genomic data, called eQTL mapping, treats gene expression profiles as quantitative traits and maps these expression traits to genomic loci (Jansen and Nap 2001; Doerge 2002; Schadt et al. 2003; Bing and Hoeschele 2005; Rockman and Kruglyak 2006; Keurentjes et al. 2007; Wang et al. 2014). Likewise, metabolomic expression profiles can be also treated as quantitative traits and mapped to genomic loci, *i.e.*, mQTL mapping (Keurentjes et al. 2006; Schauer et al. 2006; Dumas et al. 2007; Gieger et al. 2008; Illig et al. 2010; Suhre et al. 2011; Wei et al. 2017). Both eQTL mapping and mQTL mapping are derivatives of QTL mapping. Genes and metabolites that are mapped to the same loci as a trait may be used to uncover the biological networks that govern the variability of the trait. Moreover, combining the additional omic datasets with genomic data in selection analysis has potential to improve trait predictability.

Various omic datasets have been used for prediction of the EBVs of agronomic traits. For example, transcriptomic data have been used to predict hybrid performance (Stokes et al. 2010; Fu et al. 2012), and transcriptome-based prediction in hybrid maize appeared to be more precise than genome-based prediction (Frisch et al. 2010). Similarly, genomic data and metabolomic data of two backcross populations from 359 recombinant inbred lines (RILs) were used to predict biomass of Arabidopsis thaliana (Gärtner et al. 2009), in which the predictabilities for two prediction strategies were very close,*i.e.*, 0.17 and 0.16 for genomic prediction and metabolomic prediction, respectively. A population was generated by testcrossing 285 diverse Dent inbred lines from worldwide sources with two testers and used to predict the combining ability for seven biomass- and bioenergy-related traits (Riedelsheimer et al. 2012). The average predictabilities of these seven traits for genomic prediction and metabolomic prediction were 0.54 and 0.33, respectively. A three-step prediction strategy was proposed and evaluated using a wheat dataset which consists of 1,604 hybrids and their 135 parents (Zhao et al. 2015). Their results showed that for hybrids without parental line in common, hybrids sharing one parental line, and hybrids sharing both parental lines, the genome-based prediction accuracies were 0.32, 0.65 and 0.89, respectively. Note the prediction accuracy, which is a different measure from predictability, was defined as the correlation between the predicted and the observed phenotypes divided by the square root of heritability. The corresponding metabolome-based prediction accuracies were 0.15, 0.42 and 0.74, respectively.

With the explosion of omic data, how to appropriately use these resources to aid selection has become a heated topic. It has been indicated that inclusion of metabolomic data did not improve predictive value, but hampered the performance of genomic selection in hybrid wheat (Zhao et al. 2015). Prediction based on all available omic data (genomic, metabolomics and transcriptomic data) rarely outperformed the best single omic data prediction in hybrid rice when various prediction models were compared (Xu et al. 2016). However, selection by combining transcriptomic data with genomic data resulted in a higher prediction accuracy than genomic selection in maize if the omic data (genomic, metabolomic and transcriptomic data) were collected from parental lines at their early developmental stages (Westhues et al. 2017). The conflicting conclusions in the literature highlighted the need for further investigation on what combination of the omic datasets and what prediction model yields the best prediction for a trait. The answer to this question will benefit academic research and will also greatly reduce the operative cost for the industry which specializes in breeding and selection.

The goal of the study is to prove the concept that trait predictability may be optimized by using superior prediction models and selective omic datasets. For demonstration, we used an immortalized F2 (IMF2) population which was created by randomly paring 210 RILs (Hua et al. 2003). Three individual omic datasets, *i.e.*, genomic dataset, transcriptomic dataset and metabolomic dataset, and all possible combinations of these omic datasets were comprehensively analyzed for trait predictability using six widely adopted prediction methods.

## Results

### Analysis of variance for predictabilities

We calculated 168 (4×7×6) predictabilities for 4 traits using all 7 possible combinations of omic datasets (G, M, T, G + M, G + T, M + T, and G + M + T) with 6 prediction methods (Table S1; Table S2). The predictability (168 values) was treated as the response variable, and 4 traits, 7 combinations of omics datasets and 6 methods were treated as factor variables in an ANOVA analysis to detect the differences between selection schemes with different levels of these factors. The results for the IMF2 population (Table 1) show that all main and three interaction effects are significant. Comparisons between various omic data combinations with ‘method factor’ being averaged out are depicted in Figure 1. For YIELD (1^st^ panel of Figure 1), the seven combinations are classified into three levels, *i.e.*, A (best), B and C (worst). Combining genomic data and metabolomic data (G + M) produced the best predictability, while GS (prediction solely based on genomic data) gave the worst predictability. For the other three traits (KGW, GRAIN and TILLER), only two levels were detected for the seven combinations of omic datasets, with G + M being the best for KGW and GRAIN and G + M + T being the best for TILLER. Comparisons between six prediction methods with ‘combination factor’ being averaged out are depicted in Figure 2. BLUP appears to be the optimal method across all traits. For YIELD, LASSO generated the highest predictability; however, there is no statistical difference between BLUP and LASSO.

**Figure 1.**
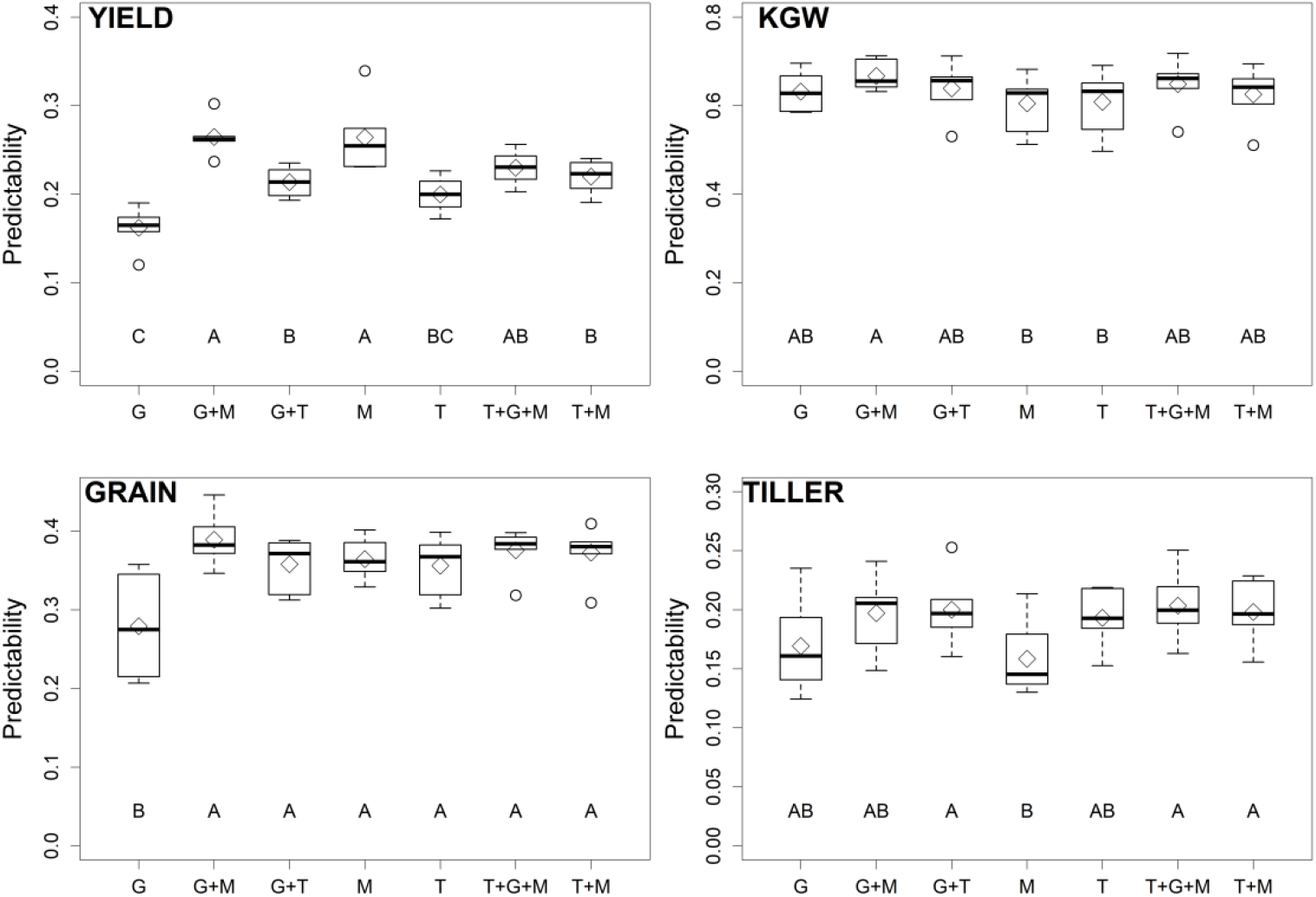
Multiple comparisons of the means of predictabilities of the four traits (YIELD, KGW, GRAIN, and TILLER) for in IMF2 population by seven combinations of omic datasets, with the differences of six prediction methods being averaged out. The capital letters ‘A’ through ‘C’ below box-plots represent the groups with significant differences in comparisons. For example, G + M (A) prediction is significantly better than G + T prediction (B), but T + G + M prediction (AB) is not significantly different from either of the other two predictions when YIELD is considered.

**Figure 2.**
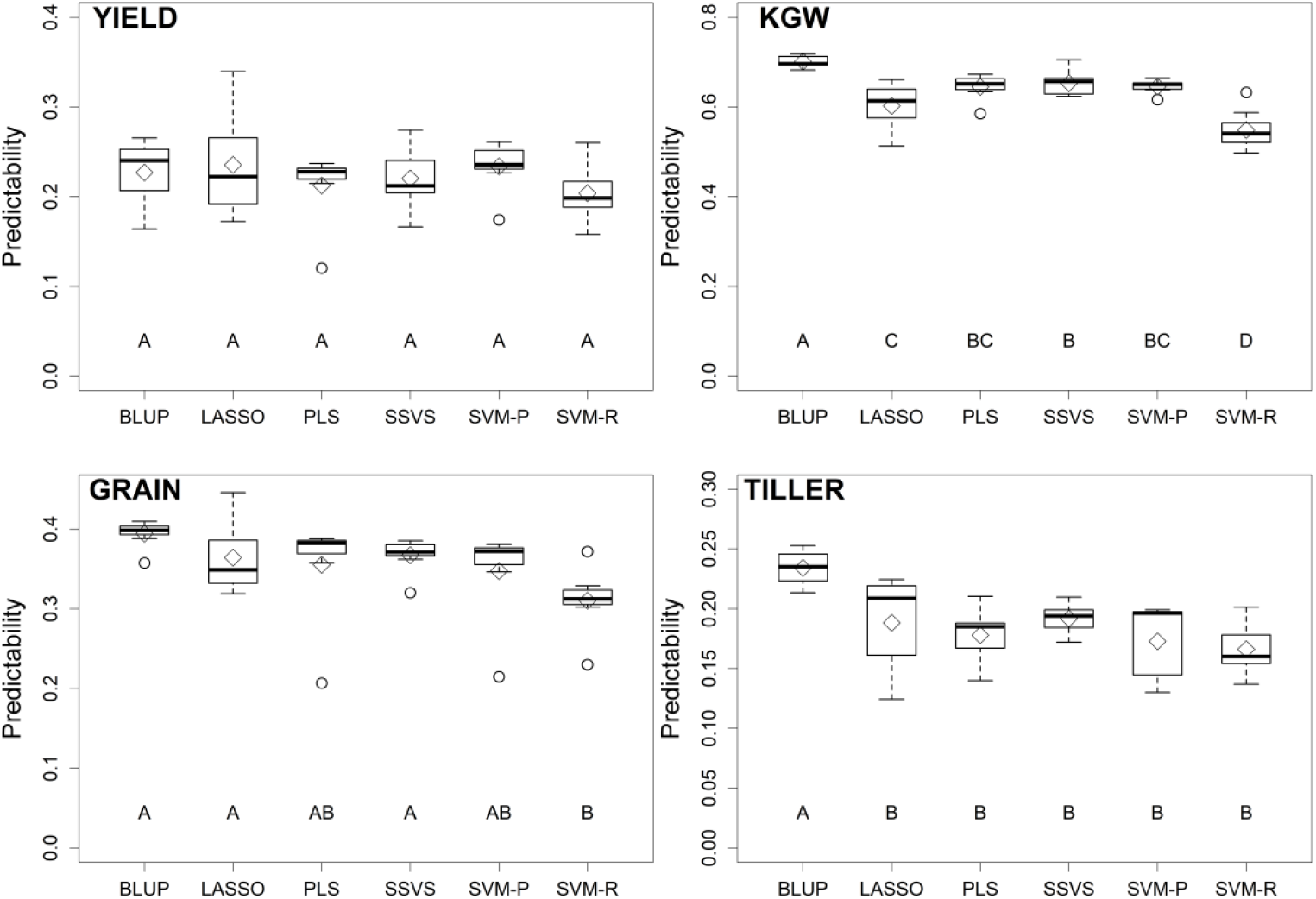
Multiple comparisons of the means of predictabilities of the four traits in the IMF2 population by six prediction methods, with the differences between seven combinations of omic datasets being averaged out.

**Table 1.**
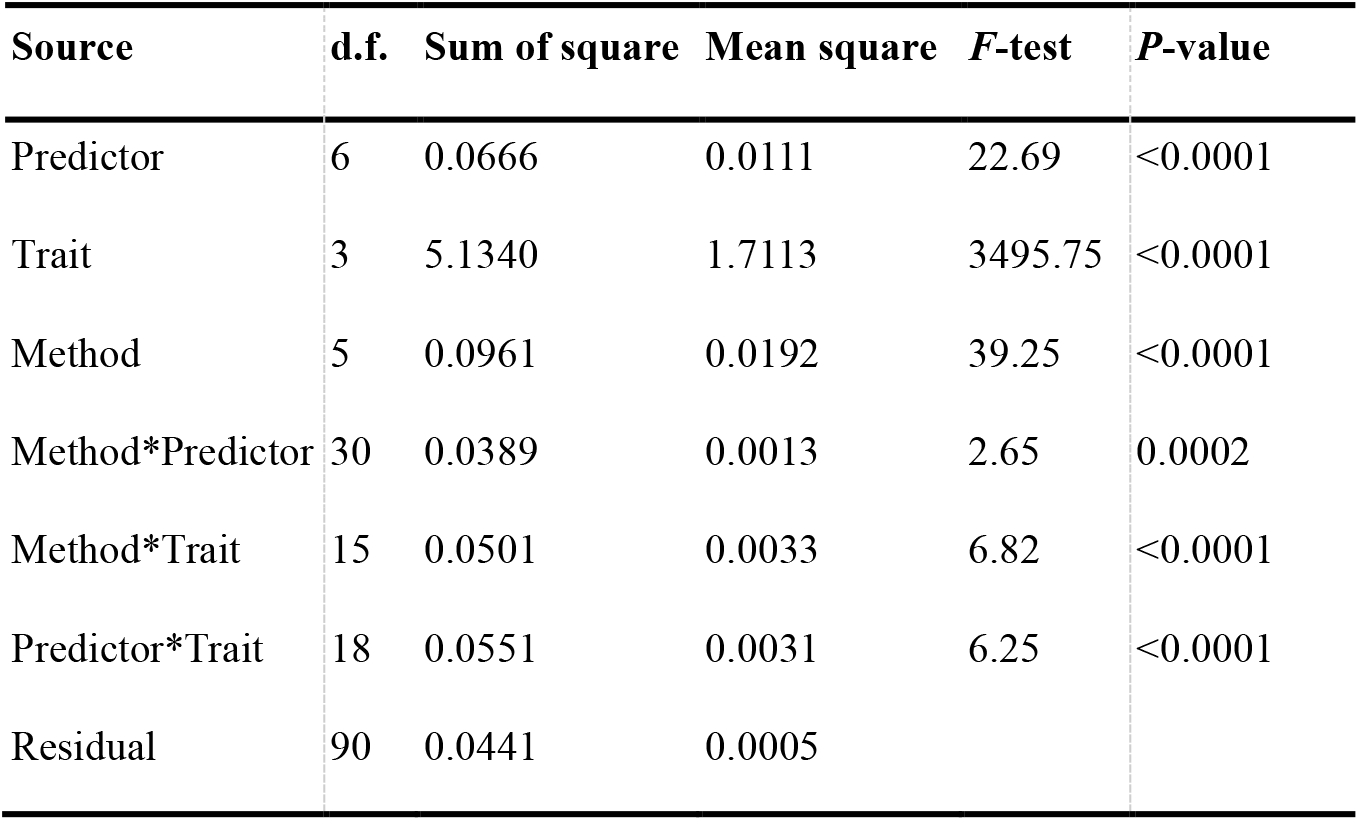
Analysis of variance of predictabilities for a IMF2 population using a 7 × 4 × 6 factorial design (seven combinations of omic datasets, four traits, and six prediction methods)

Similar analyses have been performed on the RIL population. All main and interaction effects are significant in RILs (Table S3). Comparisons between various omic data combinations with ‘method factor’ being averaged out suggest that G + M is the best prediction scheme for YIELD, KGW and GRAIN. For TILLER, the best predictability was achieved by using genomic data G only; however, the difference between G + M and G is not significant (Figure S1). BLUP outcompeted other prediction methods again in the analysis of the RIL population (Figure S2).

### Effects of different variables under different models

We calculated the effects of variables included in different models (G, M, T, G + M, G + T, M + T, and G + M + T) for 4 traits with the BLUP method since it appeared to be the optimal prediction method in both populations. All predictors (variables), including 1619 genomic variables, 1000 metabolites, and 24,994 transcripts, had been standardized before this analysis. Comparisons of the estimated effects between various models for the IMF2 and the RIL populations are depicted in Figure 3 and Figure S3, respectively. The results suggested that estimated effects of genomic and metabolomic variables are generally larger than those of the transcriptomic variables. Also, the effects of each type of omic variables under the combined model (G + M + T) are lower than those in the models where single omic data was used. In addition, the distribution of the effects of the genomic variables and metabolomic variables under the fully combined model (G + M + T) is similar with that of the G+M model.

**Figure 3.**
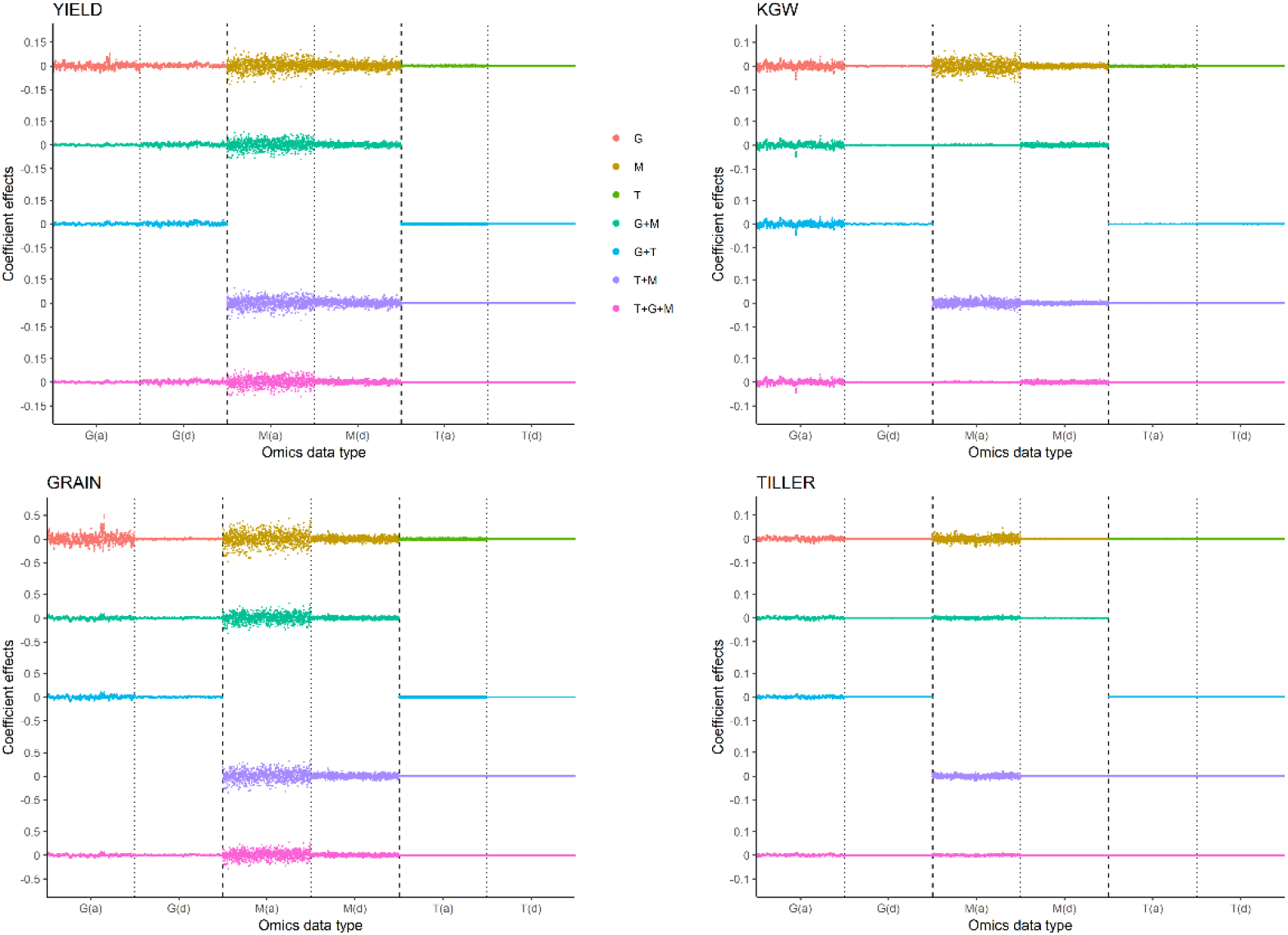
Coefficient effects with different omic datasets for the four traits in the IMF2 population. The dashed lines separate various omic-specific variables, with G, M, and T representing genomic, metabolomic, and transcriptomic variables, respectively. The dotted lines separate the additive (a) and dominance (d) variables within single omic-type variables.

### Computational efficiency

We evaluated the computational efficiency (in terms of computing time in hours) across various omic combinations and prediction methods on a regular personal computer (Intel Core i7 CPU 7700K, 4.20 GHz, Memory 16.00G). For both IMF2 population (Table S4) and RIL population (Table S5), we observed that BLUP achieved the greatest computational efficiency in average. Moreover, the computing time for BLUP increased modestly as the number of predictors increased when compared to the other methods.

### Heritability vs. predictability

The values of overall heritability of the four traits (YIELD, KGW, GRAIN and TILLER) in two populations (IMF2 and RIL) were previously calculated (Xu et al. 2016) and used in our study. The predictabilities for these four traits in the IMF2 population (average across all methods and omics combinations) were 0.2211, 0.6187, 0.3488 and 0.1794, respectively. The correlation between the heritability and the predictability for these four traits was 0.9603 ( *P* = 0.040) in the IMF2 population. Similarly, the predictabilities for these four traits in the RIL population were 0.4260, 0.6807, 0.5259 and 0.3828, respectively, and the correlation between heritability and predictability was 0.9440 ( *P* = 0.040). As expected, trait predictability generally increases with trait heritability.

### Overfitting

The squared correlation between the observed trait values and the predicted EBVs is called goodness of fit if no cross validation is applied, which is different from how predictability is defined. The measure of overfitting is the difference between the square root of goodness of fit and the square root of predictability. This is equivalent to the calculation of difference between the two correlation coefficients, one calculated between the observed trait values *vs*. the predicted EBVs without cross validation and the other one calculated with cross validation (Heslot et al. 2012). The levels of overfitting in the analyses of hybrids using various omic data combinations and prediction methods are listed in Figure 4, Figure 5 and Table S6. BLUP and LASSO were overall least affected by overfitting compared to the other prediction methods (Figure 5; Table S6); the difference between BLUP and LASSO is not statistically significant. Figure 4 suggested that G + M scheme is overall least affected by overfitting. Regarding the trait TN, G was visually less affected by overfitting than G + M; however, no statistical difference has been detected between the G and the G + M models.

**Figure 4.**
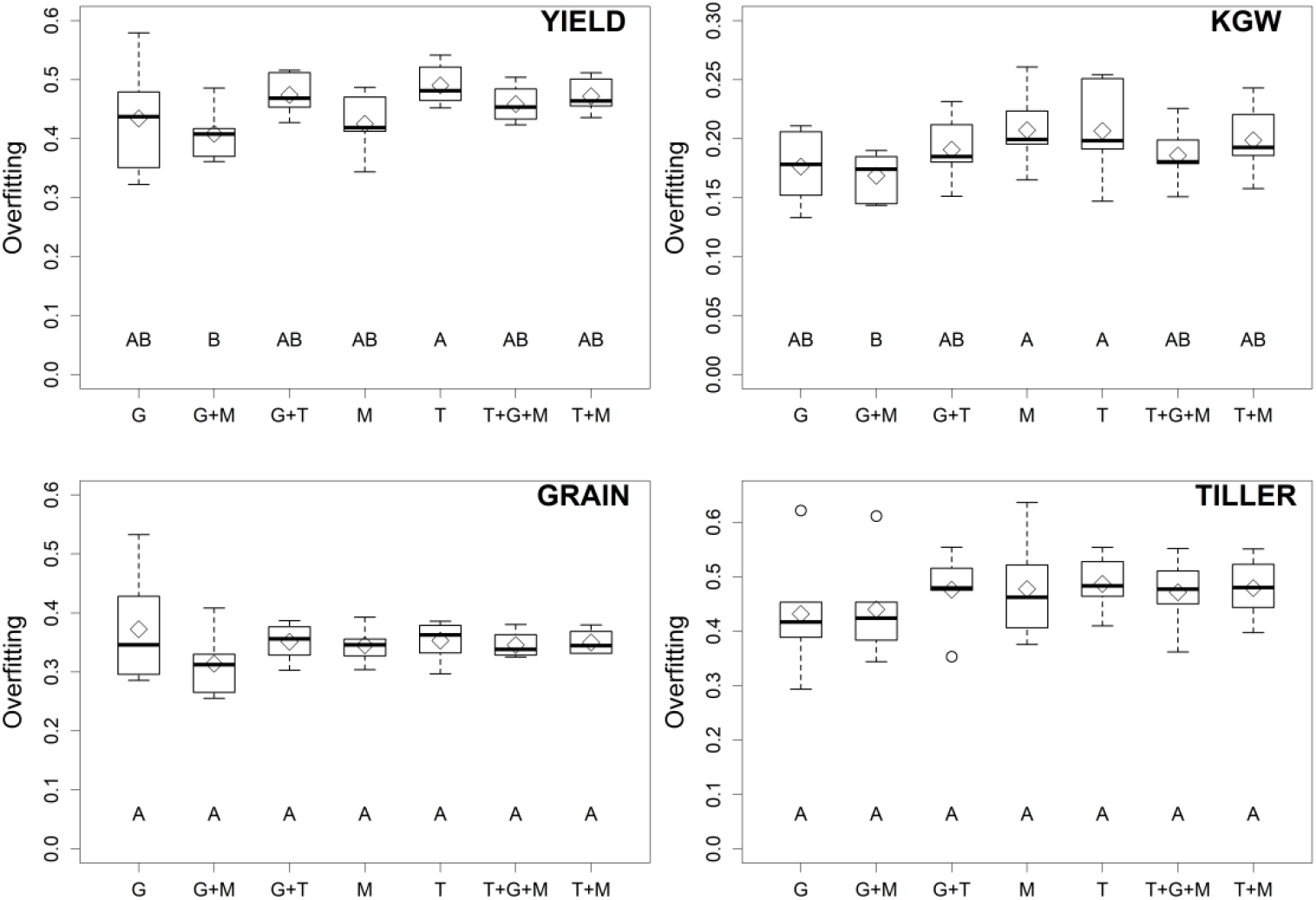
Multiple comparisons of the means of levels of overfitting for the four traits in the IMF2 population by the seven combinations of omic datasets, with the differences between the six prediction methods being averaged out.

**Figure 5.**
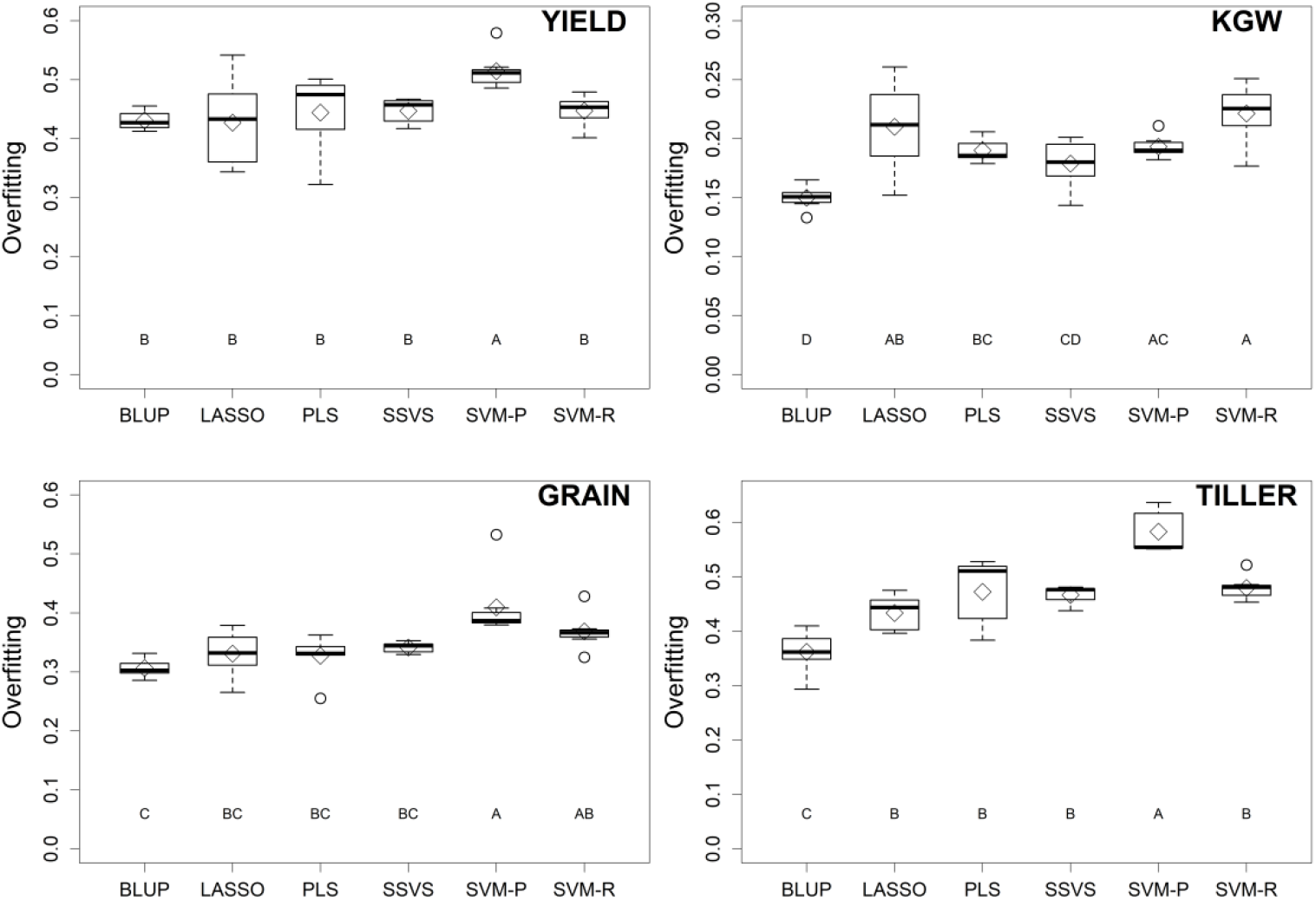
Multiple comparisons of the means of levels of overfitting for the four traits in the IMF2 population by the six prediction methods, with the differences between the seven combinations of omic datasets being averaged out.

### Selection of top crosses

The 278 experimental hybrids only represent a small subset of all 21945 possible crosses that could have been produced by the 210 RILs. For each trait, we therefore used the parameters estimated from the training samples (278 hybrids) to make predictions for all 21945 crosses. The 21945 possible crosses were then sorted based on the phenotypic values (from largest to smallest) predicted using different omic data combinations or different prediction methods. Example Data S1 shows the predicted phenotypic values of all 21945 hybrids with the BLUP method using all possible combinations of the omic data. Top 10 hybrids of each sorted list are compared in two ways since we conclude the optimal strategy for predicting hybrid rice is the BLUP method using the G + M model: (1) we first compared the top 10 hybrids selected by 6 prediction methods using G + M, and then (2) compared the top 10 hybrids selected by BLUP when different omic data combinations were used in regression. In comparison (1), out of the top 10 hybrids selected using BLUP, 9, 3, 6 and 7 hybrids were also selected by at least one other prediction method for four traits (YIELD, KGW, GRAIN and TILLER), respectively (Table S7). In comparison (2), out of the top 10 hybrids selected with G + M, 10, 8, 10 and 9 hybrids were also selected by at least one other omic data combination for four traits, respectively (Table S8).

## Discussions

This is the first study that systematically compares various trait prediction schemes using all possible combinations of omic datasets with different prediction models in order to identify the optimal strategy to achieve the best predictability. We found that the prediction based on the combination of genomic data and metabolomic data (G + M) produces the best result in the IMF2 rice population. Moreover, genomic prediction (G) or metabolomic prediction (M) is generally more effective than transcriptomic prediction (T). Inclusion of transcriptomic data to genomic prediction, metabolomic prediction, or prediction based on G + M impairs the overall model performance rather than increase predictive value. It is likely because transcriptome does not provide more information for the trait than the sum of genome and metabolome. Rather, the computational complexity is substantially increased when including transcriptomic data in the models because the number of predictor variables becomes much larger. The majority of transcripts included in the prediction models are irrelevant to the trait, leading to severe overfitting and therefore reduced predictability in cross validation. Considering YIELD, the greatest predictability was achieved by using metabolomic data (M) with LASSO, suggesting an optimal prediction strategy for prediction of yield of hybrid rice. In the RIL population, the combination of genomic data and metabolomic data (G + M) appeared to be a better option. We conclude that transcriptomic data is not necessary for selection of rice, which may greatly reduce labor and cost in industry and in future research. We also observed that the predictabilities for RILs were generally higher than those in hybrids, especially for predictions using metabolomic and transcriptomic data. This might be due to the fact that the metabolomic and transcriptomic data were directly measured for RILs but indirectly inferred, potentially with errors, for hybrids from the RIL parents. The predictabilities for hybrids may be improved if either metabolomic data or transcriptomic data or both are directly measured from the hybrids.

The effects of genomic and metabolomic variables under different models are generally larger than those of the transcriptomic variables. Moreover, the effects of the transcriptomic variables are generally lower than those of the genomic variables and the metabolomic variables in the G + M + T model. The sum of these evidences confirmed the reliability of using G + M model in hybrid rice selection. We also noticed that the effects of genomic variables and metabolomic variables in the G + M model were both smaller than their counterparts in the G model or M model where single omic datatype was analyzed. This result indicated that genomic data and metabolomic data provide very similar information for prediction of traits, and, therefore, when included in the same model (G + M), their effects were compromised compared to the single-omic-data models (G or M). However, the increased predictability in G + M model compared with the single-omic-data models (G or M) justified the use of the combination of genomic data and metabolomic data in hybrid rice selection. In addition, the effects of the genomic and metabolomic variables under the G + M + T model are very similar to that of the G + M model, which supported our argument that transcriptomic data is not necessary in rice selection when genomic and metabolomic data are available.

BLUP appeared to be a robust prediction method since the variation of the BLUP predictabilities of various omic data combination is small compared to those for the other prediction methods. Note that the computing time of BLUP depends on the number of kinship matrix rather than the number of variables used for calculation of the kinship matrices. Whereas, the computing time of the other five prediction methods substantially increases with the number of variables in the models. The number of kinship matrices (covariance structures) used in BLUP for the hybrid population is twice as many as that for the RIL population; nevertheless, this does not significantly increase the total computational time. The much higher trait predictabilities achieved by the BLUP method made this method more desirable than other methods.

Among the six prediction methods, SVM-POLY has the greatest goodness of fit (Table S9); however, the predictability of SVM-POLY is unfavorable. This suggests that goodness of fit is not suitable for evaluating prediction models and the potential overfitting may undermine the predictive value. Rather, the predictability, which is equivalent to the square of the difference between the square root of goodness of fit and the level of overfitting, can objectively reflect the applicability of the models when they are applied to independent datasets rather than training set. In our rice study, BLUP appeared to have the highest predictabilities and lowest levels of overfitting in hybrids (Table S1; Table S9; Figure 5), indicating that BLUP is more efficient in capturing signal from noise than the other prediction methods.

We also examined the prediction performance for four traits based on the data in years 1998 and 1999, respectively, using the BLUP method with various combinations of omic datasets. It seemed that the predictabilities for individual years were lower than that can be achieved with the combined data (averaged trait values across years) (Figure S4), indicating possible environmental variability in different years. Inclusion of environmental factor and its interaction with omic datasets may produce better trait predictabilities than simply averaging the trait values across years.

The best individuals, for example top 10 in a population, predicted by each method are often compared to see how many are in common such that the reliability of the method of interest can be evaluated. Considering G + M, an average of 6.3 top hybrids (out of top 10) selected by the BLUP were also selected by at least one of other five methods. In addition, an average of 9.5 top hybrids (out of top 10) selected with G + M model were also selected by at least one other omic data combinations when the BLUP was applied. These results further confirmed the reliability of our selection model using the BLUP method with the G + M combination.

For YIELD, the predictabilities for BLUP, SSVS and SVM-POLY were close to each other. Among the top 10 hybrids selected by the BLUP, 7 were selected by SSVS and 6 were selected by SVM-POLY. It appeared that methods with similar predictabilities tend to select more common top individuals. For KGW, the predictability for BLUP was significantly higher than other methods; thus, less common top hybrids are expected between BLUP and other methods. Indeed, only 3 out of the top 10 hybrids selected by BLUP were also selected by at least one other method. For GRAIN, 6 out of the top 10 hybrids selected by BLUP were also selected by at least one other method. For TILLER, BLUP achieved the highest predictability. The method with the second highest predictability was PLS which shared 4 common best hybrids with BLUP, and this number was larger than the number of common top hybrids shared by BLUP and other methods. The G + M model, of which the predictability was higher than that of G model and M model, shared an average of 6.3 top hybrids with G model only and another average of 6.3 top hybrids with M model only, and with about 4 common hybrids selected by all three models (G+ M, G and M). The results indicated that genomic data and metabolomic data contribute overlapping and complementary information on traits and the model utilizing both data, *e.g.*, the G + M model, benefits trait prediction most.

The current study has provided a guideline for rice selection in terms of what types of omic datasets and what prediction model should be used to achieve the greatest predictability. The answer may vary when different traits are considered. For other types of crops, such as maize and wheat, similar studies may be conducted to develop a selection guideline for industry practice or scientific research.

## Methods

### Rice data

Shanyou 63, an elite hybrid that has been widely cultivated in the last three decades in China, was derived from the cross between Zhenshan 97 and Minghui 63. A total of 210 RILs were derived by single-seed descent from this hybrid. An ^“^immortalized F2^”^ (IMF2) population was derived from randomly crossing these 210 RILs (Hua et al. 2002; Hua et al. 2003). Field data of four traits were considered, including yield (YIELD), 1000 grain weight (KGW), number of grains per panicle (GRAIN) and number of tillers per plant (TILLER). For the RIL population, each trait was measured from four replicated experiments (1997 and 1998 from one location, 1998 and 1999 from another location). In each replicated experiment, eight plants were sampled from each line and the average trait value was treated as the phenotypic value for this line in this experiment (Xing et al. 2002; Yu et al. 2011). For the IMF2 population, eight plants from each random cross were sampled and the average trait value was used as the phenotypic value for the F2 progeny of that cross. Trait values for each cross were measured twice in two consecutive years (1998 and 1999).

Three omic datasets, *i.e.*, genomic dataset, transcriptomic dataset, and metabolomic dataset, were only collected from the 210 RILs. Xie et al. (2010) and Yu et al. (2011) derived an ultra-high-density linkage map for these RILs, yielding genotype data represented by 1619 genetic bins. For each RIL, a genetic bin takes genotype value of 1 if the DNA in this bin is from Zhenshan 97, and 0 from Minghui 63. The transcriptomic data consisted of 24,994 gene expression traits measured in tissues sampled from flag leaves of the 210 RILs in 2008 (Wang et al. 2014). RNAs were extracted from two biological replicates of each line, and then mixed in a 1:1 ratio for expression profiling by microarrays. Robust multi-array average (RMA) analysis was used for background correction and normalization. The metabolomic data for the 210 RILs consisted of 683 metabolites measured from flag leaves and 317 metabolites measured from germinated seeds (Gong et al. 2013). Two biological replicates were sampled for flag leaves in 2009, while for germinated seeds one biological replicate was sampled in 2009 and the second biological replicate was sampled in 2010. Metabolomic data in both tissues were log2-transformed for statistical analysis to meet with the normality assumption. The average of two replicate measurements for a metabolite was used for analysis.

The genotype of an IMF2 hybrid was deduced from the genotypes of two crossing parents. Let 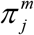 and 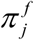 be *p*×1 vectors of the genotypes (1 for Zhenshan 97 and 0 for Minghui 63) for male and female RIL parents, respectively, where *m* = 1619. We define additive genotype of the IMF2 individual as

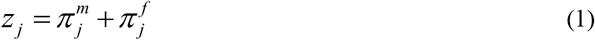

and dominance genotype as

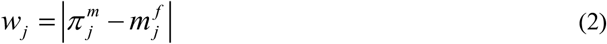

with *j* = 1,…, *q*, where *q* = 278. Therefore, the additive genotypes for the IMF2 population is defined as

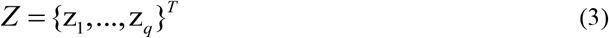

and the dominance genotypes for the IMF2 population is defined as

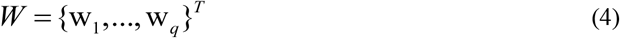

For the IMF2 population,

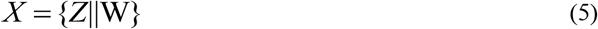

is a *q*× *2p* genotype matrix. Likewise, the metabolomic and transcriptomic data for the IMF2 population were not directly measured; rather, they were calculated from two crossing parents of each IMF2 hybrid
in a similar way, with 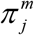 and 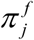 representing metabolomic or transcriptomic measurements for the two RIL patents.

### Prediction methods

Six statistical methods were used for prediction: (i) LASSO developed by (Tibshirani 1996) and implemented by GlmNet R program (Friedman et al. 2010); (ii) Henderson’s BLUP implemented in the R program written by (Xu et al. 2016); (iii) SSVS (also called Bayes B) developed by (George and McCulloch 1993); (iv) support vector machine using the radial basis function (SVM-RBF) implemented in the R package kernlab (Karatzoglou et al. 2004); (v) support vector machine using the polynomial kernel function (SVP-POLY) implemented in the R package kernlab (Karatzoglou et al. 2004); and (vi) partial least squares (PLS) implemented in the R package pls (Wehrens and Mevik 2007).

For the linear methods (LASSO, BLUP, SSVS and PLS), the single-omic-data regression is

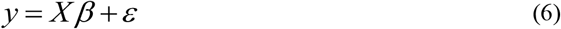

where *y* is the trait values, predictor variables *X* may be one of *X_SNP_*, *X_MET_* and *X_EXP_*, where *SNP, MET* and *EXP* indicate the three omic datatypes, β is the vector of regression coefficients, and ε is the random error which is normally distributed with N(0, σ^2^). The fully combined-omic-data regression becomes

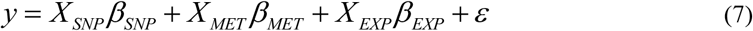

whereas other omic-data combined models have reduced format. Note in the BLUP method, more than one kinship matrix is needed to handle the mutually independent omic datasets. For IMF2 population with fully combined-omic-data regression, six kinships matrices were included in the model, with one for the additive effects and the other one for the dominance effects for each omic datatype.

Kernel methods are a class of algorithms for pattern recognition in machine learning. The most commonly used kernel methods include support vector machine (SVM) in which various kernel functions may be used for describe the relationship between dependent variable *y* and explanatory variable *X*, i.e.,

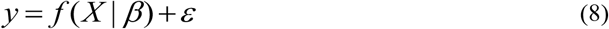

Where

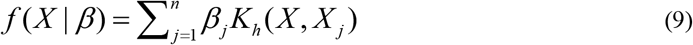

and *K_h_* (*X, X_j_*) is a kernel selected. In this study, we chose the Gaussian kernel (SVM-RBF) and the polynomial kernel (SVM-POLY) for implementation of SVM functions.

### Cross-validation

In this study, a 10-fold cross-validation was used to evaluate the predictability of each prediction method and combination of omic datasets. The trait predictability is defined as the squared correlation between the observed trait values and the predicted EBVs in cross-validation environment. The predictability calculated for a sample depends on how the sample is partitioned into different subsets for cross-validation. Therefore, 100 repeated cross-validations were performed for each analysis by randomly partitioning data in different ways and the average of the 100 predictabilities from the 100 repeated cross-validations was used for the study.

